# Closely related *Bacteroides* of the murine intestinal microbiota affect each other’s growth positively or negatively

**DOI:** 10.1101/2023.03.05.530569

**Authors:** Hanna Fokt, Gabija Sakalyte, Rahul Unni, Mohammad Abukhalaf, Liam Cassidy, Georgios Marinos, Maxime Godfroid, Birhanu M Kinfu, Ruth A Schmitz, Christoph Kaleta, Andreas Tholey, John F Baines, Tal Dagan, Daniel Unterweger

## Abstract

The mammalian intestine is a unique ecosystem for thousands of bacterial species and strains. How naturally coexisting bacteria of the microbiota interact with each other is not yet fully understood. Here, we isolated formerly coexisting, closely related strains of the genus *Bacteroides* from the intestines of healthy, wild-derived mice. The effect of one strain on another strain’s growth was tested in 169 pairs *in vitro*. We find a vast diversity of growth promoting and growth inhibiting activities. A strong positive effect was observed between two strains with differing metabolisms. Growth inhibition among a subset of strains was associated with the known bacterial toxin bacteroidetocin B. Across all strains, we observed growth promotion more often than growth inhibition. The effects were independent of two strains belonging to the same or different species. In some cases, one species differed in its effect on another according to host origin. These findings on obligate host-associated bacteria demonstrate that closely related and naturally coexisting strains have the potential to affect each other’s growth positively or negatively. These results have implications for our basic understanding of host-associated microbes and the design of synthetic microbial communities.

## Introduction

Members of the intestinal microbiota live in a complex ecosystem. The mammalian intestine harbours thousands of diverse microbes in close association with the host and with direct access to the host’s diet. Changes in community composition are associated with diseases ranging from chronic inflammatory disorders to obesity (1, 2), indicating the importance of ecological factors that affect the growth and survival of bacteria. The genus *Bacteroides* is of particular interest, as it is one of most prominent members of the most abundant phyla in the mammalian intestine, *Bacteroidota*. Among many other health-related phenomena, they metabolise polysaccharides, and are generally associated with a healthy microbiota (3–5). Most attempts to understand how ecology affects the growth and survival of *Bacteroides* species have focused on the host and its diet (6–9). Another factor could be the microbial community a bacterium lives in and the interactions between the bacteria therein.

A thorough understanding of host-associated bacterial communities requires knowledge of how bacteria affect each other. Bacteria-bacteria interactions can be positive, negative, or neutral. Existing theory provides a framework that explains the evolution of mutualistic and competitive interactions between bacteria (10). Mathematical modelling predicts that community functions, such as metabolic breakdown, and features, such as stability and resilience, depend on interactions between community members (11–15). While theoretical and model-based approaches are commonly utilized, comparatively few experiments have been performed to test these hypotheses, leaving room for controversy (16). Existing studies on synthetic communities, which mostly comprise strains with different origins that do not interact in nature, provide a glimpse of the complex interaction networks in bacterial communities (17–20). Depending on the size and phylogenetic diversity within their respective communities, individual strains often represent an entire species, genus, or phylum, enabling comparisons of interactions at high taxonomic levels like the bacterial order (18). More studies on natural populations and naturally co-existing bacteria at lower taxonomic levels are needed to provide a more representative understanding of the number and kind of bacteria-bacteria interactions in the microbiota.

Existing experimental work on the molecular mechanisms of bacteria-bacteria interactions gives insight into the diverse tools with which bacteria can interact with each other (21). For example, *B. ovatus* is known to positively affect the growth of *B. vulgatus* (now also referred to as *Phocaeicola vulgatus*) by the secretion of enzymes for the digestion of inulin, thereby producing metabolites that enhance the growth of the acceptor strain (22). *B. vulgatus* itself was shown in another study to inhibit the growth of other *Bacteroides* species by producing bacteroidetocin A, a diffusible peptide belonging to a large family of bacterial toxins (23). Another toxin with antibacterial activity, Fab1, is produced by enterotoxigenic and nontoxic *B. fragilis* strains (24). Whereas these mechanisms demonstrate the capability for interactions, generalizing these results and applying this knowledge to communities of naturally co-existing strains and species remains a challenge.

Here, we perform controlled experiments in the laboratory to test the effect of bacteria on each other’s growth among natural populations of *Bacteroides*. Our focus is on closely related strains and species that have yet to be studied in depth. We isolated strains of three *Bacteroides* species, *B. acidifaciens*, *B. caecimuris*, and *B. muris* from multiple individual mice (Table S1). The isolates showed the potential to affect each other’s growth either positively or negatively. Some of these phenotypes are associated with toxic proteins and differences in metabolism. Interestingly, we also find that strains of naturally co-existing bacterial species from one host individual affect each other differently than isolates of the same two species from another host individual.

## Results

### Bacteroides strains are inhibited or promoted by each other’s culture supernatants

To determine how the growth of one strain is affected by the growth of another, we exposed each strain to filtered spent media (also referred to as supernatant) of all the other strains in the study. This experiment allowed us to measure the effect of the changes in media caused by the growth of one strain (nutrient depletion, degradation and production of metabolites, secretion of proteins, etc.) on the growth of another. A total of 169 strain combinations were tested, including 13 strains each belonging to one of three species, *B. acidifaciens* (n=5), *B. caecimuris* (n=6), or *B. muris* (n=2), the latter of which was recently described (25, 26). We grew the strains as monocultures in spent media of self or nonself and measured the optical density (OD) at the final time point as a read-out of bacterial growth. When comparing growth in nonself to self, we observed multiple cultures with significantly higher or lower relative OD values (Fig. 1a, Table S2). These findings suggest varying positive or negative effects of different *Bacteroides* strains on each other’s growth.

**Figure 1.**
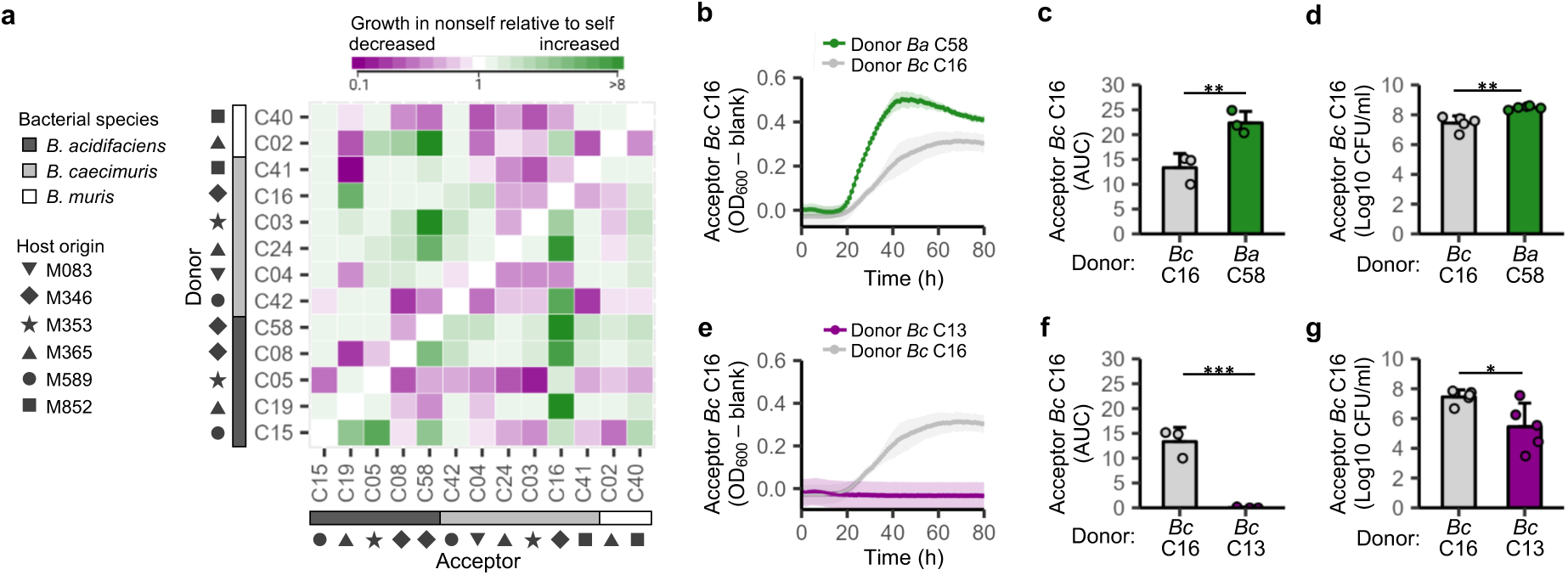
Growth inhibition and growth promotion by supernatant of *Bacteroides* strains. (a) Growth of acceptor strains in spent media of indicated donor strains relative to growth in self. Each strain (Cxx) belongs to one of three species (indicated with bars) and has been isolated from the caecum of one of six mice (indicated with symbols). Median of six independent experiments is shown. (b, c, e, f) Growth curve and corresponding area under the curve (AUC) measurements of the acceptor strain in spent media of the indicated donor strains. For the AUC data, mean and standard deviation of three independent experiments (individual dots) are shown. The presented growth curves show the mean and standard deviation of two cultures per treatment of one of the three experiments. (d, g) Counts of colony-forming units (CFU) of the acceptor strain after growth in spent media of the indicated donor strain. Mean and standard deviation of five independent experiments (individual dots) are shown. Significant differences were determined by the Wilcoxon test. * *p* < 0.05, ** *p* < 0.01, *** *p* < 0.001.

To validate our observations, we focused on strain *Bc* C16 and measured its growth in self and in spent media of strain *Ba* C58, which had a strong positive effect in the screen. When exposed to the supernatant of *Ba* C58, *Bc* C16 has a higher growth rate and grows to a higher maximal OD than in self (Fig. 1b). Accordingly, the area under the curve (AUC) is higher and colony-forming unit (CFU) counts show a higher number of bacteria in nonself than in self (Fig. 1c, d), indicating that the supernatant of *Ba* C58 positively affects the growth of *Bc* C16. This confirms our findings on this interaction in the screen. The same strain, *Bc* C16, shows hardly any growth when exposed to the supernatant of another strain, *Bc* C13, which had been noted to engage in inhibitory activity, although it was not part of our systematic screen (Fig. 1e). The low AUC values and reduction in CFUs (Fig. 1f, g) further support our finding that *Bc* C13 negatively affects the growth of *Bc* C16.

Taken together, we find that different *Bacteroides* strains affect each other’s growth in different directions, whereby the growth of one isolate could be promoted or inhibited by exposing it to the supernatants of different isolates. Next, we explored potential molecular mechanisms underlying these observations.

### Ba C58 and Bc C16 differ in their metabolism

Strain *Bc* C16 stood out in the screen by its increased growth as an acceptor in spent media of multiple strains. We hypothesized that the positive effect on the growth of *Bc* C16 could be mediated by differences between the metabolism of the donor and acceptor strains. Metabolic modelling based on whole genome sequences predicted that multiple strains (*Ba* C05, *Ba* C08, *Ba* C15, *Ba* C19, *Ba* C58) all shared 100% of their metabolic pathways, while other strains showed differences in the metabolic pathways they possessed (Fig. S1). The observation that strains with identical metabolic pathways produced supernatants with differing growth promoting activities demonstrates that metabolic modelling based solely on the presence/absence of genes is insufficient to predict all effects on growth. *Bc* C16, which grows better in the supernatant of *Ba* C58 (Fig. 1b), was predicted to differ in its metabolism from the latter (Fig. 2a). While *Ba* C58 and *Bc* C16 share most metabolic pathways, as expected from two strains of the same genus, we detected pathways of multiple metabolic classes like chemoautotrophic energy metabolism and UDP-sugar biosynthesis that are present in *Ba* C58, but absent from *Bc* C16 (Fig. S2). To test the prediction of the model experimentally, we measured the metabolic activity of these strains in the presence of 95 metabolites. We found a small but statistically significant difference in the metabolism of the two strains (Fig. 2b). Such subtle metabolic differences could result in the metabolic by-products of one strain promoting the growth of another strain that is not able to generate these products itself. A transfer of metabolites from *Ba* C58 to *Bc* C16 was predicted by a modified metabolic model with customized parameters according to the conditions of our experimental set-up (Fig. S3, Table S3).

**Fig. 2.**
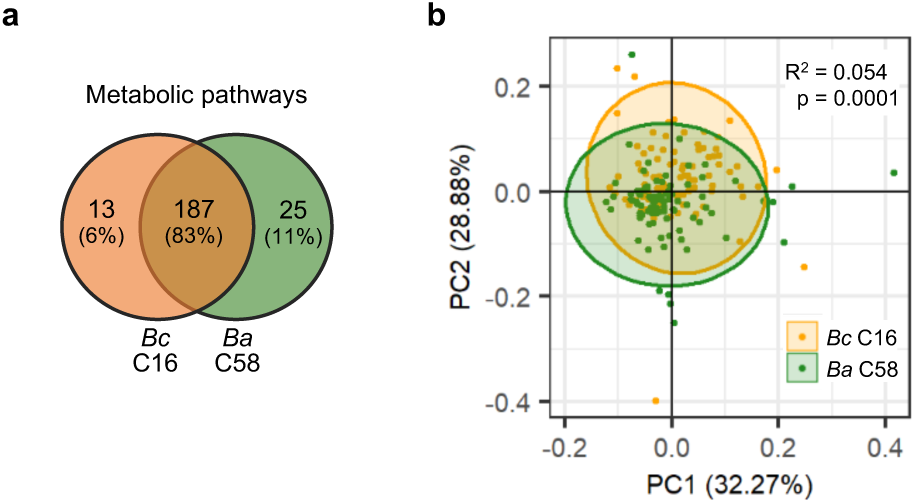
Two strains Bc C16 and Ba C58 differ in their metabolism. (a) Predicted presence and absence of metabolic pathways based on whole genome analysis. (b) PCA plot of each strain’s growth in 95 different metabolites (individual dots). Statistical analysis was performed with Adonis (10000 permutations). The analysis was performed on the data of four independent experiments.

Thus, in sum, we observed subtle metabolic differences between two strains, one of which grows better in the supernatant of the other.

### Inhibiting supernatant of strains Ba C05 and Bc C13 contains anti-bacterial proteins

To better understand the molecular mechanism underlying growth inhibition, we focused on two strains, *Ba* C05 and *Bc* C13, and tested the inhibiting activity of their supernatants in a soft-agar overlay assay that is widely used to study interactions involving bacterial toxins (27). We found inhibition halos in plates embedded with *Bc* C03 after the addition of the supernatant from *Ba* C05 and *Bc* C13 (Fig. 3a). No inhibition halo was observed when supernatant was added to self, as expected from bacterial toxins that are often encoded with an antitoxin or immunity protein. Whereas the *Bc* C13 supernatant inhibits *Ba* C05, the *Ba* C05 supernatant does not inhibit *Bc* C13, suggesting that the supernatants of the two strains possess different inhibiting activities.

**Figure 3.**
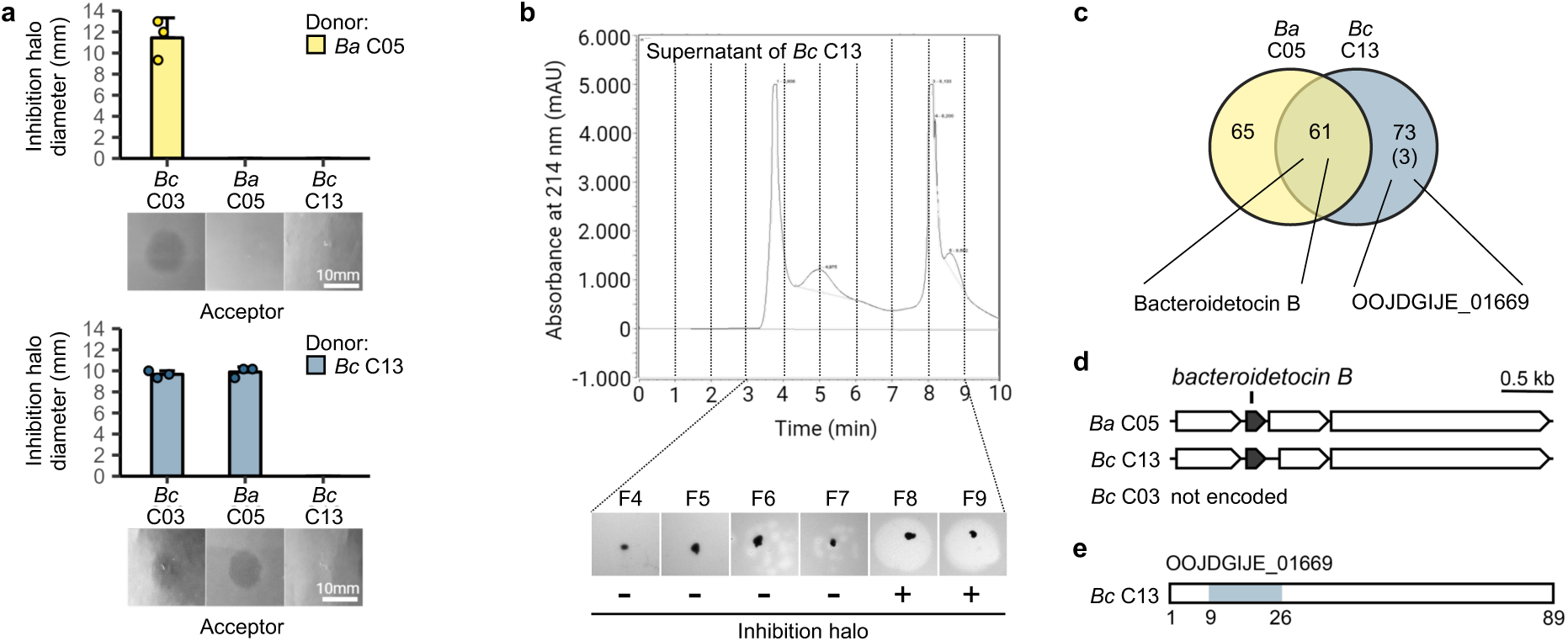
Inhibiting supernatant contains bacteroidetocin B. (a) Soft agar overlay assay with the acceptor embedded in the agar and supernatant of the donor added. Inhibition halos show inhibitory potential of *Ba* C05 and *Bc* C13. The mean and standard deviation of three independent experiments is shown. (b) Size exclusion chromatography (SEC) of supernatant of the strain *Bc* C13 and soft agar overlay assay of 6 fractions (F4-F9) to identify active fractions with inhibitory activity. Only fractions corresponding to 1-10 min collection are shown (UV absorbance at λ=214 nm). Strain *Bc* C03 was used as acceptor. (c) Venn diagram showing shared and unique proteins identified by LC-MS in SEC-fractions with inhibitory activity for the strains *Ba* C05 and *Bc* C13. The number in brackets indicates protein families of *Bc* C13 that were unique to this strain in the MS data and the comparative genome analysis. (d) Depiction of genomic locus of *bacteroidetocin B* in the indicated strains. (e) Depiction of a hypothetical protein (locus tag is indicated) with homology to a known bacteriocin. The homologous region from amino acid 9-26 is highlighted in colour.

To identify proteins that could mediate the toxic activity, we fractionated the supernatants of the two strains by size exclusion chromatography (Fig. 3b). Fractions with inhibiting activity were identified in a soft-agar overlay assay, digested with trypsin, and subjected to LC-MS for protein identification. Over 100 proteins each were identified in supernatants of *Ba* C05 and *Bc* C13 (Fig. 3c, Table S4). One of them is bacteroidetocin B, which is known for its antibacterial activity across *Bacteroides* (23), and had not yet been described of the herein studied species (Fig. 3d). It is not encoded in the genome of *Bc* C03, which is sensitive to growth inhibition by *Ba* C05 and *Bc* C13 (Fig. 3a, 3e). So far, it is not fully understood how bacteria that make bacteroidetocin B protect themselves from the toxin. Following the logic of a toxin antitoxin system that, when missing, provides a strain neither with the ability to inhibit others nor the protection of self from the toxic activity, we could speculate that *Bc* C03 might be inhibited in growth by the bacteroidetocin B of *Ba* C05 and *Bc* C13. Although the bacteroidetocin B proteins of strains *Ba* C05 and *Bc* C13 have identical amino acid sequences (Fig. S4), the supernatant of *Bc* C13 inhibits more strains than *Ba* C05 (Fig. 3a). We reasoned that *Bc* C13 might make additional, potentially unknown, antibacterial toxins. By focusing on those proteins that were unique to the supernatant of *Bc* C13 and uniquely encoded by genes of this strain (Fig. 3c), we identified three hypothetical proteins with sizes of 10-30 kDa (Table S5). One of them (locus tag OOJDGIJE_01669) has a domain that shares homology with the leader sequence of Carnobacteriocin 2, a class II bacteriocin of *Carnobacterium piscicola* (28, 29) (Fig. 3e, Table S6).

In summary, we observed that growth inhibition is specific to individual strains and found at least one protein with anti-bacterial activity in the active fractions of inhibiting supernatants.

### Interactions are predicted between strains of the same or different species

Next, we predicted how two strains would interact with each other based on our experiments with spent media and investigated these interactions across the phylogeny of the strains in this study. This analysis includes multiple strains of the same species, which is a taxonomic level that is rarely the focus of studies on bacteria-bacteria interactions. As observed in the phylogenetic tree, *B. acidifaciens* and *B. caecimuris* are more closely related to each other than to *B. muris* (Fig. 4a). Of note, the intraspecific diversity is also higher among *B. caecimuris* strains than among *B. acidifaciens* strains (Fig. S5).

**Figure 4.**
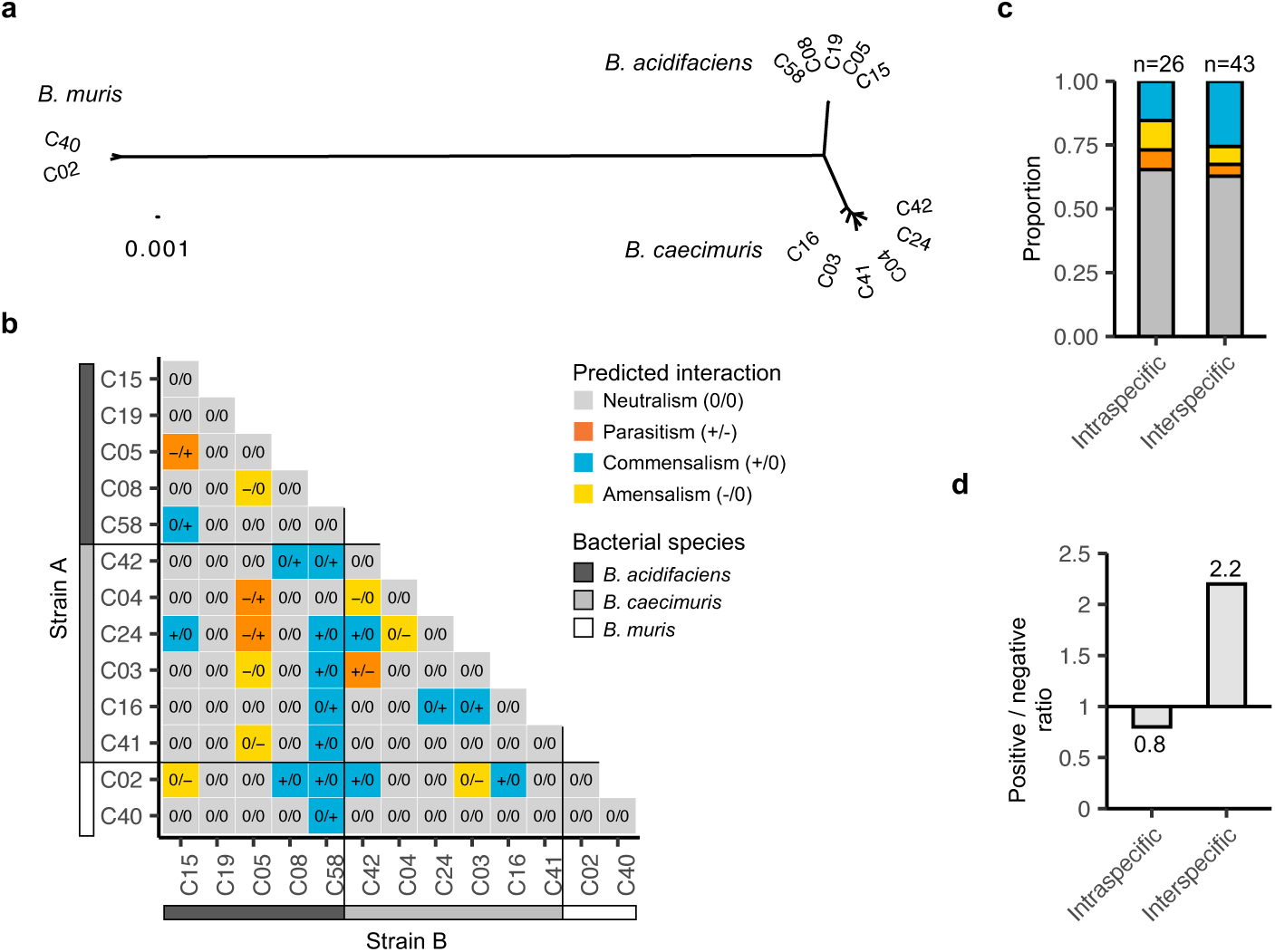
Predicted bidirectional interactions between *Bacteroides* strains. (a) Phylogenetic tree of the analysed strains based on concatenated nucleotide alignments of core genes. (b) Predicted bidirectional interactions between the indicated strains. (c) Proportion of interspecific bidirectional interactions. Statistical analysis was performed with Fisher’s Exact Test, *p* > 0.05. (d) Ratio of positive (commensalism) over negative (amensalism, parasitism) bidirectional interactions. Only interactions between strains of different host origins were considered for the analysis in (c) and (d).

To study intra- and interspecific interactions, we analysed data from the initial screen by phylogeny. We predicted bidirectional interactions between two strains based on our growth measurements of one strain in the spent medium of another strain and considered only those as interactions that had a statistically significant effect on growth (Table S2). We predicted neutral interactions (0/0), positive interactions (commensalism, +/0), and negative interactions (ammensalism, -/0; parasitism -/+) (Fig. 4b). The predicted interactions were independent of whether the strains belong to the same or different species (Fisher’s exact test, *p* = 0.6745) (Fig. 4c). This data suggests that interactions may occur between bacteria of the same species as they do between bacteria of closely related species. Within species, the ratio of positive to negative bidirectional interactions was close to one (Fig. 4d). Between species, we predicted twice as many positive bidirectional interactions as negative ones. Although this trend holds when analysing predicted unidirectional interactions only, there was no statistically significant dependence of the interaction (positive vs. negative) on the interacting strains (intraspecific vs. interspecific) (Fisher’s exact test, *p* = 0.4524) (Fig. S5). This finding suggests that the analysed strains may display a similar range of interaction, irrespective of belonging to the same or a different species.

### The effect of Ba supernatant on Bc growth varies with host origin

As the strains used in this study originated in the intestinal microbiota of multiple mice, we set out to test how strains that had co-existed naturally in the same host affected each other’s growth. We focused on *B. acidifaciens* and *B. caecimuris* strains originating from two mice, M346 and M353. Depending on the individual host, we found differences in how one species affects the other. *B. caecimuris* from mouse M353 (*Bc* C03) was inhibited when grown in spent media of *B. acidifaciens* originating from the same host (*Ba* C05). In contrast, *B. caecimuris* from M346 (*Bc* C16) showed more growth in spent media of *B. acidifaciens* from the same mouse (*Ba* C58) than in self (Fig. 5). These results show that strains of two species from one mouse affect each other in the opposite manner to strains of the same two species from another mouse.

**Figure 5.**
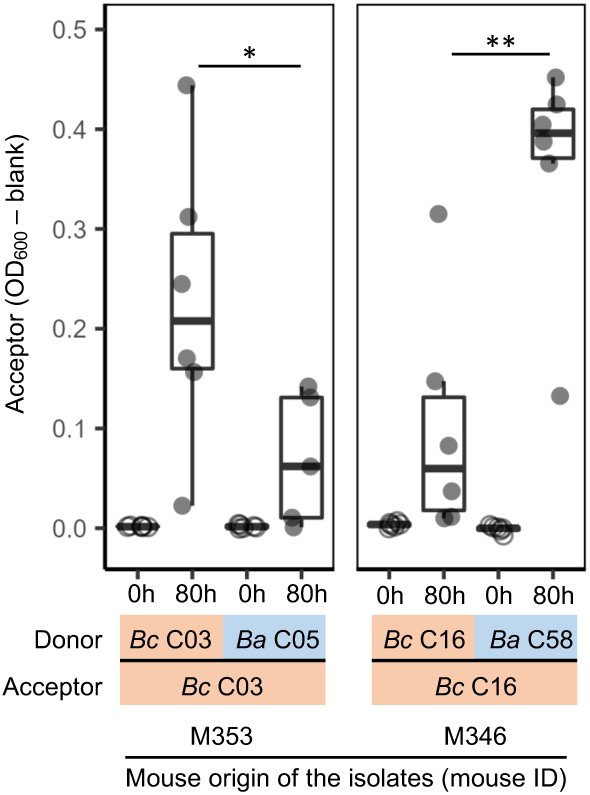
Naturally co-existing strains differ in how they affect each other’s growth based on host origin. Bacterial growth of the indicated acceptor strain in spent media of self or of *B. acidifaciens* strains that originated from the same host. Box plots are shown of six independent experiments (individual dots). Wilcoxon test was performed to test for statistical significance. *p* < 0.05 *, *p* < 0.01 **.

In light of the applied potential of tools to increase or decrease the growth of a particular species in the intestinal microbiota, we further tested supernatant of strain *Ba* C05. As this strain seemed to show a narrow spectrum of activity against non-*Bacteroides* species (Fig. S6), we wondered whether we could use it to target *Bacteroides* strains that originated from other mice. For yet unknown reasons, we did not observe growth inhibition when exposing *B. caecimuris Bc* C16 of mouse M346 to strain *Ba* C05 (Fig. S7a). However, as expected, we observed growth inhibition when exposing *B. caecimuris Bc* C24 of mouse M365 to strain *Ba* C05 (Fig. S7b).

In conclusion, we show that one species can be affected very differently by another according to the host origin and because of intraspecific diversity. Further, we demonstrate the potential and limitation to using growth inhibition between naturally coexisting strains for targeted inhibition of a strain from different host individuals.

## Discussion

In our study, we report on changes in the growth of newly isolated *Bacteroides* strains upon exposure to each other’s spent media. Strains of the three species *B. acidifaciens*, *B. caecimuris*, and *B. muris* affect each other intra- and interspecifically. As demonstrated on strains that were simultaneously isolated from the caecum of multiple hosts and were formerly coexisting, bacteria of one species grew very differently in the other species’ supernatant depending on the individual host they were isolated from. This data complements existing work by (i) focusing on closely related strains that belong to either the same or different *Bacteroides* species and (ii) emphasizing strains that naturally coexisted in the same hosts prior to isolation.

Our *in vitro* experiments captured the potential for bacteria-bacteria interactions between naturally coexisting microbes. Although the observed effects of strains on each other’s growth have not yet been tested *in vivo*, independent work has shown in general that bacteria-bacteria interactions observed in the laboratory can also be seen in a colonized host. For example, *in vitro* measurements of pairwise interactions between species of the *C. elegans* gut microbiome could predict their interactions *in vivo* (19). Further, findings on the molecular mechanisms of positive and negative bacteria-bacteria interactions among *Bacteroides* have been confirmed *in vivo* (22, 30). In comparison to *in vitro* conditions, the *in vivo* environment is not only more complex, but also subject to dynamic changes with potential effects on bacterial interactions. For example, host feeding could temporarily increase the nutrient content in the environment, which may reduce positive interactions mediated by cross-feeding (31, 32). Negative interactions may be more common in high-diversity communities, such as the *in vivo* environment, as a strain may be more likely to engage in negative interactions in the presence of other strains that potentially affect it adversely (33). Furthermore, changes in the environment along the longitudinal and horizontal axis of the gastrointestinal tract could favour one interaction over another at a given location in the gut (31).

Our data on how closely related strains affect each other’s growth complement existing data on interactions between bacteria of different orders and phyla. Whether of the same or different species, we found the bacteria to affect each other similarly. Metabolic simulations predicted that bacteria are more likely to promote each other’s growth when the difference in metabolism and phylogenetic distance are higher (34). Although we see a trend of more positive interactions between than within species, the analysed strains alone might be too closely related for this to become statistically significant.

Our results expand the network of interactions within a host and its microbes from host-microbe to host-microbe-microbe interactions. Strong indications for host-microbe interactions are derived from studies of co-evolution, which reveal genomic signatures of bacterial adaptation, possibly to the particular associated host (35). Bacteria of the Bacteroidota have repetitively been linked to the genomics of their hosts (36–38). A recent manuscript describes a genome-wide association study on the pangenome of the very same *B. acidifaciens* strains included in this study, and identified nine loci in the bacterial genomes that associate with the murine host subspecies (8). Interestingly, one of these loci is *tolC*, which encodes an outer membrane protein that is also known for its role in the import and export of bacterial toxins in Proteobacteria (39). Our phenotypic data on how *B. acidifaciens* strains affect each other’s growth, combined with data on their host-association provides a rare snapshot that simultaneously captures host-microbe and microbe-microbe interactions of one bacterial species in the microbiota.

## Materials and Methods

### Isolation of Bacteroides strains from mouse cecal content

Fourteen *Bacteroides* strains (Table S1) were isolated from the cecal contents of the stock collection of wild-derived outbred colonies of *Mus musculus musculus* (n=3 mice derived from Kazakhstan (KH), n=2 mice derived from Austria (WI)) and *M. m. domesticus* (n=1 mouse derived from Massif-Central, France (MC) and n=1 from Iran (AH)) maintained at the Max Planck Institute for Evolutionary Biology in Plön, Germany. Mouse husbandry and sampling were performed in accordance with the German animal welfare law (Tierschutzgesetz) (Permit V 312-72241.123-34).

Fresh cecal content was placed in pre-reduced brain-heart infusion (BHI) medium with 20% glycerol, homogenized, and stored at −80 °C until further processing. Schaedler Anaerobe KV (SKV) Selective Agar with Lysed Horse Blood (Thermo Scientific, Waltham, MA, US) plates were purchased ready-to-use and stored at 4 °C. Prior to inoculation, the SKV medium was reduced by placing the plates overnight under anaerobic conditions at room temperature. All the steps were performed under anaerobic conditions (gas mixture: 5% H_2_, 5% CO_2_, 90% N_2_) in a vinyl anaerobic chamber (Coy Lab Products). Cecal contents were homogenized by vortexing, and serial 10-fold dilutions in anaerobic BHI medium were made. Next, 50 μl of the last three dilutions (10^-6^, 10^-5^, and 10^-4^) were plated on SKV plates. The plates were incubated at 37 °C for 48 h. If the growth was insufficient after 48 h of incubation, the plates were incubated for an additional 24 h.

Colonies were picked from agar plates according to their morphology: *Bacteroides* form circular, white or beige, shiny, smooth colonies that are approximately 2-3 mm in diameter. Next, *Bacteroides* genus-specific primers Bac 32F and Bac 708R (40) were used for the amplification of an approx. 750 bp portion of the 16S rRNA gene by colony PCR. PCR reactions contained 3.6 μl H_2_O, 5 μl of Multiplex PCR Master Mix (Qiagen), 0.2 μl of 2 μM primers and 1 μl DNA template. Cycling conditions were as follows: initial denaturation for 15 min at 95 °C; 35 cycles of 30 s at 94 °C, 90 s at 58 °C, and 90 s at 72 °C; followed by 10 min at 72 °C and ∞ 12 °C. Colonies were confirmed to belong to *Bacteroides* sp. by Sanger sequencing and matching to the RDP database training set 14 (41).

### Genome sequencing and processing

Genomic DNA for whole-genome sequencing was extracted using the DNeasy UltraClean Microbial Kit (Qiagen). Bacterial biomass from strains grown on SKV agar plates was resuspended directly in 300 μl of PowerBead Solution (Qiagen) and vortexed to mix. All following steps were performed according to the manufacturer’s instructions. DNA samples were prepared according to Illumina Nextera XT protocol. The final DNA library was supplemented with 1% PhiX and sequenced on an Illumina NextSeq 500 system using NextSeq 500/550 High Output Kit v2.5 with 300 cycles.

Sequences were quality checked with FastQC (42) and trimmed with Trimmomatic (43) using a 4-base wide sliding window approach, cutting when the average quality per base dropped below 15. Reads with a minimum length of 40 bases were kept. Reads were assembled into contigs using SPADES (44). The quality of the assemblies was assessed with QUAST (45). The assemblies were annotated by Prokka (46).

### Homologous protein identification and clustering into families

To identify homologous proteins, all-against-all local alignments of annotated genomes were performed using the blastp v2.5.0+ (47) with an e-value threshold of 1e-5. Every pair of significant hits was aligned using the global alignment tool, powerneedle from the EMBOSS package (48, 49). Significant hits sharing at least 30% global amino acid identity were retrieved and clustered into homologous families using MCL v14-137 (50).

### Strains, culture media and growth conditions

Fourteen *Bacteroides* strains were stored as frozen monoculture stocks. Chopped meat (CM) medium containing 20 g/l meet extract, 30 g/l tryptone, 5 g/l yeast extract, 5 g/l K_2_HPO_4_, 5

µg/ml hemin, 2.5 µl/l Vitamin K1 and 0.5 g/l cysteine-HCl was used to grow the bacteria. All strains were grown at 37 °C under anaerobic conditions (gas mixture: 5% H_2_, 5% CO_2_, 90% N_2_) in a vinyl anaerobic chamber (Coy Lab Products).

### Production of bacterial spent media

*Bacteroides* strains were grown in monocultures for 96 h at 37 °C in 5 ml CM medium under anaerobic conditions. Bacterial spent media supernatants (SM) were generated by centrifugation of the grown culture at 5800 x *g* for 20 min and subsequent filter-sterilization (0.2 µm, VWR™). SM samples were prepared freshly before each experiment under anaerobic conditions. For the heat and cold susceptibility assays, SM were incubated at 99 °C for 15 min, and at -20 °C until complete freezing, respectively.

### Growth assay in bacterial spent media

Growth assay was performed in pairwise manner where each strain was used as acceptor and as a donor. The growth in self SM was used as the reference for each combination. The SM obtained as described above were supplemented by CM medium (2/3 CM medium and 1/3 SM) and inoculated with fresh bacterial cultures toa final OD_600_ of 0.01 (total volume of 200 μl). The monocultures were grown in respective SM under anaerobic atmosphere in 96-well plates (Sarstedt) for 80 h. Bacterial growth was measured using Sunrise^TM^ plate reader (TECAN, Männedorf, Switzerland). Absorption was measured at wavelength 600 nm. During continuous measurements, the plate was heated inside the reader to 37 °C and a 10 s shaking step was performed prior to each measurement. Colony forming units (CFU) were counted at 24 h of growth and area under the curve was calculated after 80 h of growth for the acceptor grown in donor’s SM and in self SM. All the experiments were performed at least three times.

The effect of one strain on another strain’s growth was determined by calculating the medians of the growth ratios as follows: OD_600_ 80 h (growth in other SM)/OD_600_ 80 h (growth in self SM), from six independent experiments. The effects of non-self SM were classified as growth promoting or growth inhibiting if the ratio were higher or lower than 1, respectively. All the OD_600_ values at 80h growth lower than respective OD_600_ at 0 h growth, were discarded.

### Soft agar overlay assay

The inhibitory potential of *Bacteroides* strains was assayed by the soft agar overlay technique (27). The OD_600_ of all bacterial cultures grown in 5 ml of CM medium was measured, the spent medium was prepared as described above. In order to normalize for the amount of the antimicrobial molecule secreted to the medium, an OD_600_ = 0.5 was chosen as a reference. Hence, 500 µl of the cultures with an OD_600_ = 0.5 were spun down to collect supernatant. For the OD_600_ above or below 0.5, the culture volume to collect was calculated accordingly.

The antimicrobial activity of cell-free spent media was assessed on lawns of each tested strain in a pairwise manner (all-against-all, including self as a control). Bacterial cultures were carefully mixed with 13 ml of soft agar 0.5% (w/v) in order to get a final OD_600_ of 0.05, and poured onto square Greiner CM agar plates 12×12 cm (Merck). Next, 5 µl of cell-free supernatant were spotted onto the lawn of solidified bacteria-soft agar mixture and allowed to dry completely. A 5 µl drop of the pure medium used as a control was spotted on each plate. The plates were incubated at 37 °C for 48 h. Bacterial growth inhibition was quantified by measuring the average diameter (in mm) of the growth inhibition halos, measured in three different points each. The experiments were performed in triplicate.

### Biolog assay

The ability of B*acteroides* strains to use different carbon sources was evaluated by performing a series of assays using Biolog AN MicroPlates™ (Biolog, Hayward, CA), following the protocol provided by the manufacturer, with minor modifications. The strains were grown in CM medium under anaerobic conditions at 37 °C for 72 h. Bacterial biomass was collected by centrifugation at 5000 x*g* for 10 min and inoculated in AN Inoculating Fluid (Biolog, Hayward, CA) such that an OD_600_ of 0.05 was achieved. The Inoculating Fluid was then pipetted into Biolog AN MicroPlates™ (100 μl per well), and the initial OD_590_ was measured using a Spark® microplate reader (TECAN, Männedorf, Switzerland). The plates were then incubated in anaerobic jars at 37 °C for 48 h, after which the final OD_590_ was measured. The difference between the final and initial OD_590_, ΔT, was used to determine the extent to which each of the metabolites were utilized by the strains. The metabolites differentially utilized by the strains were identified by testing whether the difference in ΔT for each metabolite was significantly different between each pair of strains using the Kruskal–Wallis test.

### Sample preparation and fractionation by Size Exclusion Chromatography (SEC)

Strains *Bc* C13 and *Ba* C05 were grown anaerobically on CM agar plates as described above. Bacteria and associated inhibitory molecules were carefully swabbed from the surface of the agar plates and resuspended in the PBS buffer (Carl Roth). To remove cell-associated inhibitory molecules, bacteria were vortexed for one minute and pelleted by centrifugation. The supernatants were collected, filter-sterilized (0.2 µm, VWR™) and tested for inhibitory activity using soft agar overlay assay.

Supernatants showing biological activity were concentrated using Amicon Ultra-0.5 centrifugal filters (3 kDa) (Merck KGaA, Darmstadt, Germany) as follows. Filters were washed first with 500 µl MilliQ water and centrifuged at 14,000 x*g* for 10 min. Flow-through and retentate were discarded, then 500 µl sample were added and centrifuged at 14,000 x*g* for 30 min at 4 °C; this procedure was repeated once to load all the sample volume. Filters were transferred upside down in new collection tubes and retentates were collected by centrifugation at 1,000 x*g* for 2 min, 4 °C. Samples were then fractionated on Dionex U3000 nanoHPLC system (Dreieich, Germany) equipped with BioSep SEC-S 2000 column (4.6 x 300 mm) (Phenomenex, California, US) and UV detector (λ= 214 nm) for a total run of 15 min with an isocratic flow of 0.5 ml/min (30% Acetonitrile (ACN), 0.1% trifluoroacetic acid (TFA)). Fractions were collected in *ca*. 1 min windows around the peaks and collected fractions were split equally in two (one for bioassay and one for LC-MS) and lyophilized. All the fractions (15 fractions for *Bc* C13 and 6 fractions for *Ba* C05 strain) were tested for the inhibitory activity using soft agar overlay assay.

### Sample preparation for Liquid Chromatography-Mass Spectrometry (LC-MS)

Fractions of SEC that showed inhibitory activity in the soft agar assay (fraction 8 for *Bc* C13 and fraction 5 for *Ba* C05 strain) were dissolved in 100 µl 100 mM Triethylammonium bicarbonate (TEAB), pH 8. Reduction was performed with dithiothreitol (DTT) (10 mM final concentration) for 30 min at 56 °C, followed by alkylation with Chloroacetamide (CAA) (50 mM final concentration) for 20 min at 23 °C in the dark. Proteins were digested overnight with 0.4 µg trypsin (Promega, WI, USA) in 100 mM TEAB. Subsequently, peptides were dried in a vacuum concentrator (Concentrator Plus, Eppendorf).

### Liquid chromatography–mass spectrometry

Dried peptides were resuspended in 3% ACN plus 0.1% TFA, then separated employing a Dionex U3000 nanoHPLC system equipped with an Acclaim pepmap100 C18 column (2 µm,

75 µm x 500 mm). The eluents used were; eluent A: 0.05% Formic acid (FA), and eluent B: 80% ACN + 0.04% FA. A programmed 60-minute run at 300 nl/min was performed as follows: 3 min at 4% B, 30 min linear gradient from 4% to 50% B, 1 min linear gradient from 50% to 90% B, 10 min isocratic 90% B followed by equilibration for 15 min at 4% B. Eluted peptides were analysed online on a QExactive Plus or an Orbitrap Fusion Lumos (with high-field asymmetric waveform ion mobility spectrometry (FAIMS)) mass spectrometer (Thermo, Bremen, Germany). In the QExactive Plus, a full MS scan (300-1500 m/z, resolution 70,000, AGC 3e6, Max IT 100 ms) was followed by HCD fragmentation and MS/MS scans of the 10 most intense ions (NCE 27, resolution 17,500, AGC 1e5, Max IT 50 ms, 20 s dynamic exclusion). In the Orbitrap Fusion Lumos, a full scan MS acquisition was performed (300- 1500 m/z, resolution 60,000, AGC 4e5, Max IT 100 ms) at two FAIMS compensation voltages (CV), -45 and -60 V. Subsequent data-dependent MS/MS scans were collected within a 1.5 s cycle time for the most intense ions via HCD activation at NCE 30 (resolution 15,000, AGC 5e4, Max IT 22 ms, 60 s dynamic exclusion (after 2 hits)).

### MS data analysis

MS raw data were analysed against a predicted proteome from a sequenced *B. caecimuris Bc* C13 (3,936 sequences) or *B. acidifaciens Ba* C05 (3,844 sequences) and known contaminants (cRAP) using Sequest HT search engine linked to Proteome Discoverer^TM^ 2.2 (Thermo). Enzyme specificity was set to Trypsin (Full) with a tolerance of two missed cleavages. Mass tolerances of 10 ppm and 0.02 Da were set for the precursor and fragment ion mass errors, respectively. Carbamidomethylation of cysteine was set as a fixed modification. Methionine oxidation and N-terminal acetylation were set as variable modifications. Strict parsimony criteria were applied filtering peptides and proteins at an FDR < 1%. Proteins were further filtered to contain at least two identified peptides.

### Metabolic modelling

The available genomes of all the strains were utilized for the reconstruction of the genome- scale metabolic models of these organisms. The reconstruction was conducted with the software *gapseq* (version 1.2 (51)). Specifically, we firstly reconstructed functional, high quality models based on an automatically predicted medium and then we adapted them so that they can produce biomass on an *in-silico* growth medium that simulates the experimental one. Apart from the reconstruction of the models, *gapseq* exports the information of absence or presence of certain metabolic pathways (e.g., from MetaCyc (52), custom ones) for each strain.

The models were further used to infer their metabolic interactions. To this end, we used the software *MicrobiomeGS2* (version 0.1.5, www.github.com/Waschina/MicrobiomeGS2). The step-by-step procedure of simulation includes the merging of the models in one community model whose growth is maximized. The outcome enabled us to record the metabolic exchanges among the sub-models of the strains.

In parallel, we used a 2D, spatiotemporal simulation setting as an alternative to the above described. The software that was used was *BacArena* (version 1.8.2, commit fdb02bf7 (53)). To ensure that the simulation reflects growth conditions as much as possible, while keeping the needed computational resources low, we performed growth simulations in a 170×170 grid cell, stirred environment for 8 h on the *in-silico* growth medium (downsized to fmol/grid cell) in. Each simulation was replicated ten times. Regarding the simulated scenarios, firstly, we simulated the growth of 10 copies for each model alone. Then we extracted all the compounds of the final environment at the end of the simulation (i.e., dietary compounds and metabolic products of the donor strain). Then we added these compounds along with the dietary information in a new simulation with the models of the acceptor strain and we recorded which external compounds were used by the acceptor strain.

The computational environment was of the programming language R, version 4.2.1. The main background package used for simulations was *sybil*, version 2.2.0 (54). The solver was the academic version CPLEX (version 22.1.0) that is available in R through the package *cplexAPI* (version 1.4.0). Especially for *MicrobiomeGS2*, following its default settings, the solving method was from its default to “hybbaropt”. We additionally set the upper coupling constraint “cpl_u” to 1E-6, so that we make sure the sub-models interact only if they able to grow. For data processing and visualisation, we used the R packages *ggplot2* (version 3.4.1 (55)), batman (version 1.2.1.11 (56, 57)) and tidyverse (version 1.3.2 (58)).

### Phylogenetic analysis

Phylogenetic relationships among 13 *Bacteroides* isolates were determined by performing a pangenome analysis on nucleic acid sequences using Roary v.3.13.0 with default parameters (59). Draft genomes of all strains were generated. We identified 3508 core genes in 5 *B. acidifaciens* isolates, 2293 core genes in 6 *B. caecimuris* strains, and 2806 core genes in 2 *B. muris* isolates. Seventy-eight core genes were detected among all 13 *Bacteroides* isolates. To infer phylogenetic trees, we performed a partition analysis (60) on concatenated core genome alignments using IQtree software v.2.2.0.3 with model parameters GTR+F+I (61, 62). Maximum likelihood phylogenetic trees were plotted using Figtree v.1.4.4 (63). We used the R package *ape* (64) to compute the pairwise genetic distances from the core genes phylogenetic tree.

### Biofilm prevention activity of Bacteroides against test pathogens

*Bacteroides* were grown in 50 ml pre-reduced CM medium at 37 °C until late exponential growth phase. After centrifugation, cell pellets were resuspended in 2 ml of 50 mM Tris-HCl buffer (pH 7.9) and the cells were disrupted using a French pressure cell two times (at 4.135 x 10^6^ N m^-2^) followed by centrifugation at 12,000 x g at 4 °C for 30 min. Samples were then centrifuged at 12,000 x *g* and 4 °C for 30 min. Clear lysates were subsequently filtered through 10 kDa followed by 3 kDa Amicon Ultra-0.5 centrifugal filters (Merck KGaA, Darmstadt, Germany) to generate sub-cellular fractions with molecular size of larger than 10 kDa, 3 to 10 kDa, and smaller than 3 kDa. All subcellular fractions and cell-free culture supernatant (CFCS) were filter sterilized using 0.2 µm spin filter units (Amchro, Hattersheim).

Biofilm-forming model pathogens Klebsiella oxytoca M5aI (DSM 7342), Candida albicans (DSM 11225), Staphylococcus aureus (DSM 11323), S. epidermidis RP62A (DSM 28319), Escherichia coli K12 (65), Pseudomonas aeruginosa PAO1 (DSM 1707) and Stenotrophomonas maltophilia were grown overnight in 5 ml CASO Bouillon (Roth, Karlsruhe) (C. albicans was grown in YPD medium) shaking at 120 rpm. The first two microorganisms were incubated at 30 °C and the rest at 37 °C. Grown samples were diluted in the respective medium to 10^6^ cells/ml and 150 μl (per well) was transferred to 96-well round (U)-bottom microtiter plate (MTP) (Greiner Bio-One, Kremsmünster, Österreich). To each well, 15 μl of sterile size fractionated samples and cell-free culture supernatants were supplemented. Tris-HCl buffer (50 mM, pH 7.9) was added as control. At least eight technical replicates were used per each test material. The plates were sealed with adhesive gas-permeable sterile membrane (Breathe Easy, Roth, Karlsruhe) and incubated overnight shaking at 80 rpm at the appropriate temperatures. Biofilm assays were performed using crystal violet according to (65).

### Data analysis and figures

Data was analysed using R Studio (Version 4.1.3). All non-parametric comparisons were conducted using the non-parametric Wilcoxon Rank-sum test with α = 0.05. Plots were generated using the R *gglot2* package (55). Figures were partly generated using BioRender (https://biorender.com).

## Acknowledgments

We thank Dr. Sven Künzel for support with whole genome sequencing and the animal caretakers of the MPI for Evolutionary Biology. This research was supported in part through high-performance computing resources available at the Kiel University Computing Centre. This work was funded by the Deutsche Forschungsgemeinschaft (DFG, German Research Foundation)—SFB 1182—project-ID 261376515 to RAS (Subproject Z2, B2), CK and GM (Subproject A1), AT (Subproject Z3), JFB (Subproject A2, Z3), TD (Suproject B4), DU (Subproject B4). RU is funded by the International Max–Planck Research School for Evolutionary Biology (IMPRS Evol-Bio). Work in the Unterweger Lab is supported by the German Federal Ministry for Education and Research (grant 01KI2020) and the Deutsche Forschungsgemeinschaft (RU5042, EXC2167).

## Data and Code Availability Statement

The datasets generated during and/or analysed during the current study are available from the corresponding author on reasonable request. The Whole Genome Shotgun project has been deposited at DDBJ/ENA/GenBank under the accession XXXX. The version described in this paper is version XXXX. All accessions and project IDs are given in Table Sxx. LC–MS raw data files have been deposited to the ProteomeXchange Consortium (66) by the PRIDE partner repository with the dataset identifier PXDXXXXX. The codes used in the present study are accessible via xxx.

## References

1. R. Caruso, B. C. Lo, G. Núñez, Host–microbiota interactions in inflammatory bowel disease. Nat Rev Immunol 20, 411–426 (2020).

2. M. Van Hul, P. D. Cani, The gut microbiota in obesity and weight management: microbes as friends or foe? Nat Rev Endocrinol, 1–14 (2023).

3. H. M. Wexler, *Bacteroides*: the Good, the Bad, and the Nitty-Gritty. Clin Microbiol Rev 20, 593–621 (2007).

4. A. G. Wexler, A. L. Goodman, An insider’s perspective: *Bacteroides* as a window into the microbiome. Nat Microbiol 2, 1–11 (2017).

5. F. Thomas, J.-H. Hehemann, E. Rebuffet, M. Czjzek, G. Michel, Environmental and Gut *Bacteroidetes*: The Food Connection. Frontiers in Microbiology 2 (2011).

6. C. A. Thaiss, et al., Microbiota Diurnal Rhythmicity Programs Host Transcriptome Oscillations. Cell 167, 1495–1510.e12 (2016).

7. J. L. Sonnenburg, et al., Glycan foraging *in vivo* by an intestine-adapted bacterial symbiont. Science, New Series 307, 1955–1959 (2005).

8. H. Fokt, et al., Comparative genomics of novel *Bacteroides acidifaciens* isolates reveals candidates for adaptation to host subspecies in house mice. bioRxiv 2023.01.31.526425 (2023).

9. S. Doms, et al., Key features of the genetic architecture and evolution of host-microbe interactions revealed by high-resolution genetic mapping of the mucosa-associated gut microbiome in hybrid mice. eLife 11, e75419 (2022).

10. S. A. West, A. S. Griffin, A. Gardner, S. P. Diggle, Social evolution theory for microorganisms. Nat Rev Microbiol 4, 597–607 (2006).

11. K. Z. Coyte, J. Schluter, K. R. Foster, The ecology of the microbiome: Networks, competition, and stability. Science 350, 663–666 (2015).

12. K. Z. Coyte, C. Rao, S. Rakoff-Nahoum, K. R. Foster, Ecological rules for the assembly of microbiome communities. PLOS Biology 19, e3001116 (2021).

13. R. Tsoi, et al., Metabolic division of labor in microbial systems. Proceedings of the National Academy of Sciences 115, 2526–2531 (2018).

14. A. R. Pacheco, M. Moel, D. Segrè, Costless metabolic secretions as drivers of interspecies interactions in microbial ecosystems. Nat Commun 10, 103 (2019).

15. S. Magnúsdóttir, et al., Generation of genome-scale metabolic reconstructions for 773 members of the human gut microbiota. Nat Biotechnol 35, 81–89 (2017).

16. J. D. Palmer, K. R. Foster, Bacterial species rarely work together. Science 376, 581–582 (2022).

17. M. Chatzidaki-Livanis, M. J. Coyne, L. E. Comstock, An antimicrobial protein of the gut symbiont *Bacteroides fragilis* with a MACPF domain of host immune proteins. Mol Microbiol 94, 1361–1374 (2014).

18. J. Kehe, et al., Positive interactions are common among culturable bacteria. Sci Adv 7, eabi7159 (2021).

19. A. Ortiz, N. M. Vega, C. Ratzke, J. Gore, Interspecies bacterial competition regulates community assembly in the C. elegans intestine. ISME J 15, 2131–2145 (2021).

20. A. S. Weiss, et al., *In vitro* interaction network of a synthetic gut bacterial community. ISME J 16, 1095–1109 (2022).

21. L. García-Bayona, L. E. Comstock, Bacterial antagonism in host-associated microbial communities. Science 361, eaat2456 (2018).

22. S. Rakoff-Nahoum, K. R. Foster, L. E. Comstock, The evolution of cooperation within the gut microbiota. Nature 533, 255–259 (2016).

23. M. J. Coyne, et al., A family of anti-*Bacteroidales* peptide toxins wide-spread in the human gut microbiota. Nat Commun 10, 3460 (2019).

24. Y. Bao, et al., A Common Pathway for Activation of Host-Targeting and Bacteria- Targeting Toxins in Human Intestinal Bacteria. mBio 12, e0065621 (2021).

25. H. Fokt, et al., *Bacteroides muris sp. nov.* isolated from the cecum of wild-derived house mice. Arch Microbiol 204, 546 (2022).

26. A. Afrizal, et al., Enhanced cultured diversity of the mouse gut microbiota enables custom-made synthetic communities. Cell Host Microbe 30, 1630–1645.e25 (2022).

27. K. L. Hockett, D. A. Baltrus, Use of the Soft-agar Overlay Technique to Screen for Bacterially Produced Inhibitory Compounds. J Vis Exp, 55064 (2017).

28. L. E. Quadri, M. Sailer, K. L. Roy, J. C. Vederas, M. E. Stiles, Chemical and genetic characterization of bacteriocins produced by Carnobacterium piscicola LV17B. J Biol Chem 269, 12204–12211 (1994).

29. T. Sprules, K. E. Kawulka, A. C. Gibbs, D. S. Wishart, J. C. Vederas, NMR solution structure of the precursor for carnobacteriocin B2, an antimicrobial peptide from Carnobacterium piscicola. European Journal of Biochemistry 271, 1748–1756 (2004).

30. A. J. Verster, et al., The Landscape of Type VI Secretion across Human Gut Microbiomes Reveals Its Role in Community Composition. Cell Host Microbe 22, 411–419.e4 (2017).

31. A. S. Weiss, et al., Nutritional and host environments determine community ecology and keystone species in a synthetic gut bacterial community. bioRxiv 2022.11.24.516551 (2022).

32. M. Murillo-Roos, H. S. M. Abdullah, M. Debbar, N. Ueberschaar, M. T. Agler, Cross- feeding niches among commensal leaf bacteria are shaped by the interaction of strain- level diversity and resource availability. ISME J 16, 2280–2289 (2022).

33. A. L. Gould, et al., Microbiome interactions shape host fitness. Proc Natl Acad Sci U S A 115, E11951–E11960 (2018).

34. S. Giri, et al., Metabolic dissimilarity determines the establishment of cross-feeding interactions in bacteria. Curr Biol 31, 5547–5557.e6 (2021).

35. N. A. Moran, J. P. McCutcheon, A. Nakabachi, Genomics and evolution of heritable bacterial symbionts. Annu Rev Genet 42, 165–190 (2008).

36. J. Wang, et al., Analysis of intestinal microbiota in hybrid house mice reveals evolutionary divergence in a vertebrate hologenome. Nat Commun 6, 6440 (2015).

37. M. Groussin, et al., Unraveling the processes shaping mammalian gut microbiomes over evolutionary time. Nat Commun 8, 14319 (2017).

38. M. C. Rühlemann, et al., Genome-wide association study in 8,956 German individuals identifies influence of ABO histo-blood groups on gut microbiome. Nat Genet 53, 147– 155 (2021).

39. E. Cascales, et al., Colicin biology. Microbiol Mol Biol Rev 71, 158–229 (2007).

40. A. E. Bernhard, K. G. Field, Identification of nonpoint sources of fecal pollution in coastal waters by using host-specific 16S ribosomal DNA genetic markers from fecal anaerobes. Appl Environ Microbiol 66, 1587–1594 (2000).

41. J. R. Cole, et al., Ribosomal Database Project: data and tools for high throughput rRNA analysis. Nucleic Acids Research 42, D633–D642 (2014).

42. S. Andrews, FastQC: A quality control tool for high throughput sequence data. Available online at: http://www.bioinformatics.babraham.ac.uk/projects/fastqc/ (2010).

43. A. M. Bolger, M. Lohse, B. Usadel, Trimmomatic: a flexible trimmer for Illumina sequence data. Bioinformatics 30, 2114–2120 (2014).

44. A. Bankevich, et al., SPAdes: A New Genome Assembly Algorithm and Its Applications to Single-Cell Sequencing. Journal of Computational Biology 19, 455–477 (2012).

45. A. Gurevich, V. Saveliev, N. Vyahhi, G. Tesler, QUAST: quality assessment tool for genome assemblies. Bioinformatics 29, 1072–1075 (2013).

46. T. Seemann, Prokka: rapid prokaryotic genome annotation. Bioinformatics 30, 2068– 2069 (2014).

47. S. F. Altschul, W. Gish, W. Miller, E. W. Myers, D. J. Lipman, Basic local alignment search tool. J Mol Biol 215, 403–410 (1990).

48. S. B. Needleman, C. D. Wunsch, A general method applicable to the search for similarities in the amino acid sequence of two proteins. J Mol Biol 48, 443–453 (1970).

49. P. Rice, I. Longden, A. Bleasby, EMBOSS: the European Molecular Biology Open Software Suite. Trends Genet 16, 276–277 (2000).

50. A. J. Enright, S. Van Dongen, C. A. Ouzounis, An efficient algorithm for large-scale detection of protein families. Nucleic Acids Res 30, 1575–1584 (2002).

51. J. Zimmermann, C. Kaleta, S. Waschina, gapseq: informed prediction of bacterial metabolic pathways and reconstruction of accurate metabolic models. Genome Biol 22, 81 (2021).

52. R. Caspi, et al., The MetaCyc database of metabolic pathways and enzymes and the BioCyc collection of pathway/genome databases. Nucleic Acids Res 44, D471–480 (2016).

53. E. Bauer, J. Zimmermann, F. Baldini, I. Thiele, C. Kaleta, BacArena: Individual-based metabolic modeling of heterogeneous microbes in complex communities. PLoS Comput Biol 13, e1005544 (2017).

54. G. Gelius-Dietrich, A. A. Desouki, C. J. Fritzemeier, M. J. Lercher, sybil – Efficient constraint-based modelling in R. BMC Systems Biology 7, 125 (2013).

55. H. Wickham, ggplot2: Elegant graphics for data analysis. Springer-Verlag New York, ISBN 978–3-319-24277-4 (2016).

56. O. Keyes, et al., batman: Convert Categorical Representations of Logicals to Actual Logicals (2015) (March 4, 2023).

57. J. Hao, W. Astle, M. De Iorio, T. M. D. Ebbels, BATMAN—an R package for the automated quantification of metabolites from nuclear magnetic resonance spectra using a Bayesian model. Bioinformatics 28, 2088–2090 (2012).

58. H. Wickham, et al., Welcome to the Tidyverse. Journal of Open Source Software 4, 1686 (2019).

59. A. J. Page, et al., Roary: rapid large-scale prokaryote pan genome analysis. Bioinformatics 31, 3691–3693 (2015).

60. O. Chernomor, A. von Haeseler, B. Q. Minh,, Terrace Aware Data Structure for Phylogenomic Inference from Supermatrices. Systematic Biology 65, 997–1008 (2016).

61. S. Kalyaanamoorthy, B. Q. Minh, T. K. F. Wong, A. von Haeseler, L. S. Jermiin, ModelFinder: fast model selection for accurate phylogenetic estimates. Nat Methods 14, 587–589 (2017).

62. S. Tavare, Some probabilistic and statistical problems in the analysis of DNA sequences. Some mathematical questions in biology / DNA sequence analysis edited by Robert M. Miura (1986) (February 28, 2023).

63. http://tree.bio.ed.ac.uk/software/figtree/ FigTree (February 14, 2023).

64. E. Paradis, K. Schliep, ape 5.0: an environment for modern phylogenetics and evolutionary analyses in R. Bioinformatics 35, 526–528 (2019).

65. N. Weiland-Bräuer, L. Saleh, R. A. Schmitz, Functional Metagenomics as a Tool to Tap into Natural Diversity of Valuable Biotechnological Compounds. Methods Mol Biol 2555, 23–49 (2023).

66. J. A. Vizcaíno, et al., ProteomeXchange provides globally coordinated proteomics data submission and dissemination. Nat Biotechnol 32, 223–226 (2014).

