## Supplementary figures S1-7 for "Closely related *Bacteroides* of the murine intestinal microbiota affect each other’s growth positively or negatively"

Figure S1-S7

Supplementary Tables

Table S1-S6

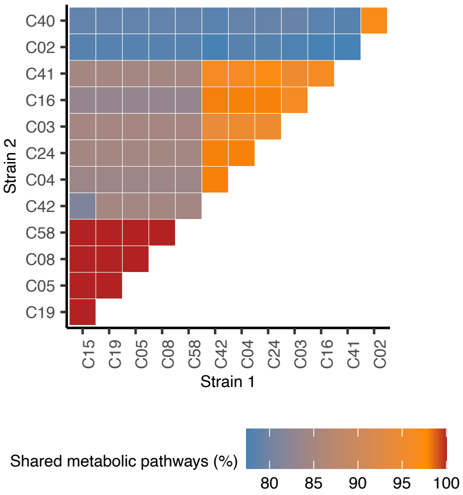


**Figure S1. Predicted shared metabolic pathways based on whole genome sequences.**

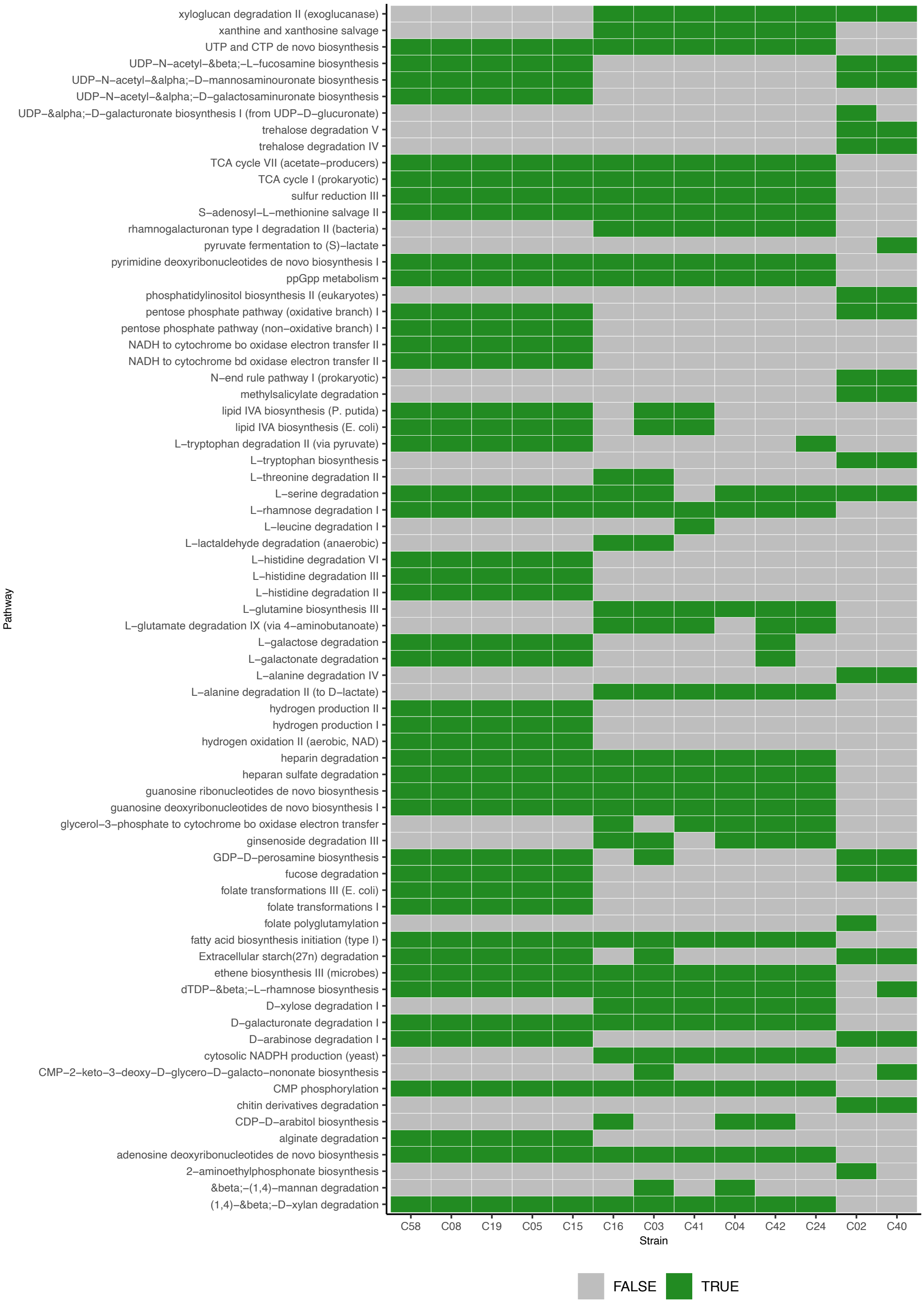


**Figure S2. Predicted presence and absence of metabolic pathways.** Only those pathways are shown that differ in presence/absence between the analysed strains.

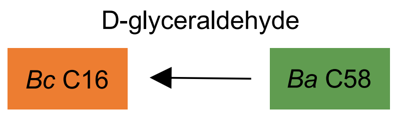


**Figure S3. Predicted transfer of metabolites from strain *Ba* C58 to *Bc* C16.**

```
>Protein alignment Alignment of 2 sequences: Bact_B_C05, Bact_B_C13

Identities = 57/57 (100%),
Positives = 57/57 (100%), Gaps = 0/57 (0%)

Bact_B_C05    1 MKKVSNKVLSRAFGGKMFGSKTECERWEGGCRCRQVYYTFWVKSYGAWNPAADWQCN 57
                MKKVSNKVLSRAFGGKMFGSKTECERWEGGCRCRQVYYTFWVKSYGAWNPAADWQCN
Bact_B_C13    1 MKKVSNKVLSRAFGGKMFGSKTECERWEGGCRCRQVYYTFWVKSYGAWNPAADWQCN 57
```

**Figure S4. Alignment of the amino acid sequences of bacteroidetocin B of the indicated strains.**

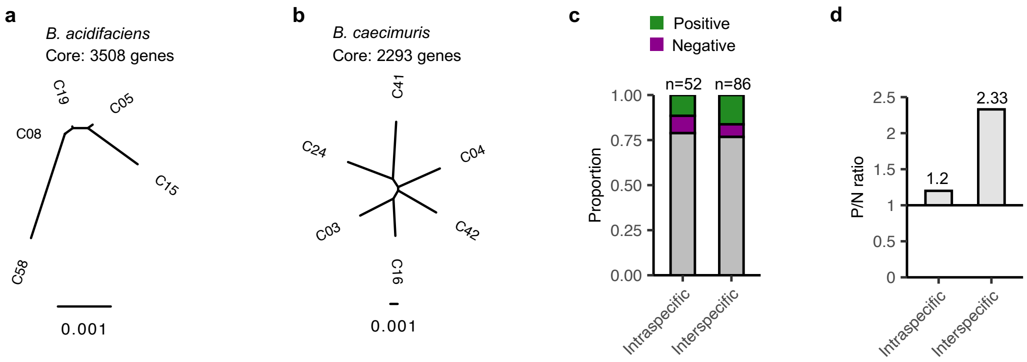


**Figure S5. Intraspecific diversity and predicted unidirectional interactions.** (a) Phylogenetic tree based on core genes of the *B. acidifaciens* isolates. (b) Phylogenetic tree based on core genes of the *B. caecimuris* isolates. (c) Proportion of predicted unidirectional positive and negative interactions. An interaction was considered positive or negative when the acceptor strain showed a statistically significant increase or decrease in growth relative to self. Statistical significance was assessed with the Fisher's exact test ($p = 0.4524$). (d) Ratio of positive (P) over negative (N) predicted unidirectional interactions.

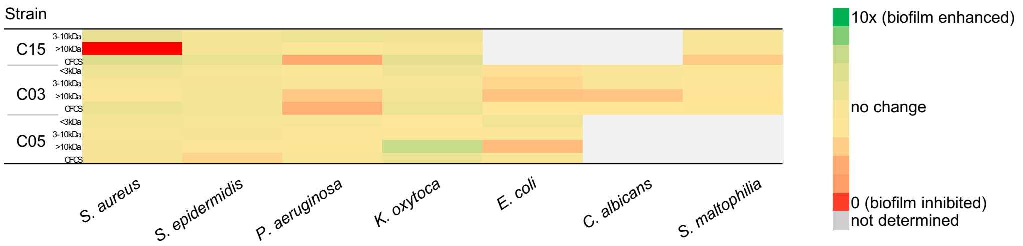


**Figure S6. *Bacteroides* strains inhibit biofilm formation of different pathogens.** Cell-free culture supernatant (CFCS) and three subcellular fractions based on molecular size (smaller than 3 kDa, between 3 and 10 kDa and bigger than 10 kDa) from each of the indicated *Bacteroides strains* were tested against *Staphylococcus aureus*, *S. epidermidis*, *Pseudomonas aeruginosa*, *Klebsiella oxytoca*, *Escherichia coli*, *Candida albicans*, and *Stenotrophomonas maltophilia.* Biofilm formation was evaluated in a crystal violet assay and inhibitory activity was calculated relative to the control (Tris-HCl buffer). At least eight technical replicates were performed.

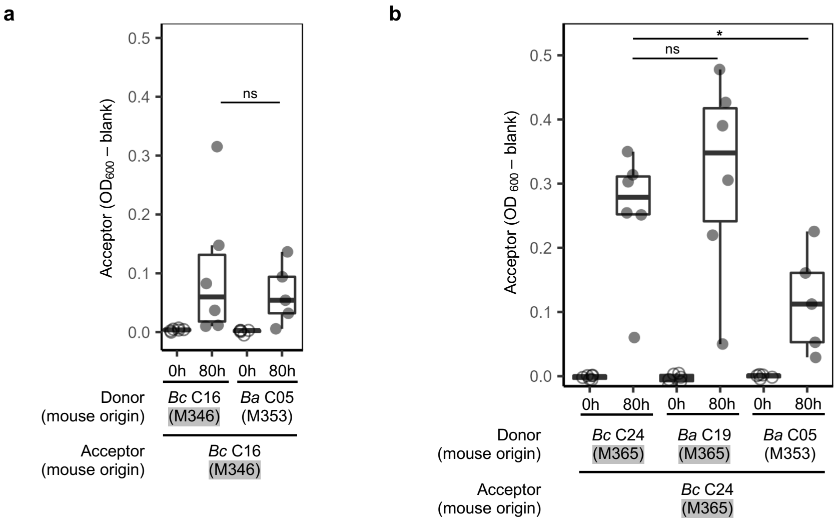


**Figure S7. Interactions between strains of the same and different host origin.** (a) Growth of *B. caecimuris* strain *Bc* C16 upon exposure to spent media of *B. acidifaciens Ba* C05 (M353). Host origins are indicated in brackets. This data was collected in the same experiment than the data shown in Fig. 5 and shows the same values for self. (b) Growth of *B. caecimuris* strain *Bc* C24 upon exposure to self, spent media of *B. acidifaciens Ba* C19, which originated from the same host, and *B. acidifaciens Ba* C05 of mouse M353. The experiments were performed six times independently. Wilcoxon test was used to test for statistical significance. ns, not significant; $p < 0.05$ *.
